# Theta phase-locked memory reactivation during REM sleep reduces memories’ emotional tone

**DOI:** 10.64898/2026.06.05.730360

**Authors:** João Patriota, Toon Brouwer, Mathijs Bergers, Timo van Hattem, Kiki Wijers, Werner Meulmeester, Elsa Juan, Umberto Olcese, Lucia Talamini

## Abstract

REM sleep appears to be involved in emotional memory processing, yet the nature and neural underpinnings of this involvement remain largely unclear. Recent findings suggest REM-related theta activity may play a central role. To test this, we developed an automated protocol to target non-arousing acoustic memory cues to specific phases of REM sleep theta oscillations. During sleep, participants who had undergone a fear conditioning procedure combined with a memory recollection and emotional evaluation task, were re-exposed to conditioned stimuli, timed to either the upswing or downswing of theta waves. Both up- and down-targeted memory cues significantly enhanced the theta dynamic. Moreover, theta phase-locked memory reactivation significantly reduced the emotional tone of conditioned memories compared to sham reactivation (no acoustic memory cue), without affecting recognition performance or dream qualities. This constitutes the first evidence that REM sleep theta oscillations are causally involved in emotional memory processing, specifically in attenuating memories’ emotional charge. These findings have important implications for understanding emotional memory consolidation and may lead to therapeutic applications in disorders featuring maladaptive emotional memories.

## Introduction

Every day, sleeping organisms enter an ecologically challenging behavioral state, during which the processing of external sensory stimuli is heavily reduced. Although this disconnection makes us vulnerable to environmental threats, the neurochemical and endocrine status of the offline brain favors a host of processes that are essential to survival ^1–6^. Besides general ‘housekeeping’ processes (e.g. energy conservation and restoration of metabolic homeostasis ^7^, these include a crucial and well known role in memory consolidation ^5,8^ and emotional regulation ^3,9–14^.

While the role of NREM sleep in declarative memory consolidation has been firmly established, with specific neural mechanisms identified ^15–18^, the contribution of rapid eye movement (REM) sleep is less clear. Findings in human subjects suggests that REM sleep may support emotional memory processing ^12–14,19–22^. Indeed, the limbic system appears relatively activated during REM sleep ^23^ while REM sleep dreams frequently feature emotional content related to daytime experience ^24,25^. Also, dreaming of aversive life events has been associated with emotional catharsis ^26^ while fragmented REM sleep has been correlated to poor emotional recovery and depressive symptoms ^27,28^. However, REM sleep’s precise role in emotional memory processing has long remained elusive, with recent reviews suggesting it may or may not strengthen emotional memories, depending on procedure, and it may either potentiate or diminish memories’ emotional charge ^11,29^.

An approach that allows direct manipulation of memory reactivation during sleep is, so-called, targeted memory reactivation (TMR). This entails biasing sleep-related memory reactivation towards specific memory items, using sensory (often acoustic) memory cues, presented during sleep ^30–32^. Applied in animal experiments, TMR has delivered strong evidence supporting memory reprocessing and consolidation during sleep ^29,31–36^. It has also been applied as a non-invasive tool to study memory consolidation in humans, providing strong evidence for the role of non-REM (NREM) sleep in memory consolidation ^29,37^. Few studies to date have used TMR during REM sleep ^38–41^ and even fewer have specifically addressed emotional memory consolidation ^42–45^. The studies that did, produced varying results across different memory parameters and paradigms ^29^.

In terms of the neural mechanisms underlying REM sleep-related memory processing, research has focused on theta oscillations (4 - 8 Hz), the predominant low frequency in the REM sleep spectrogram. REM theta oscillations reflect synchronized activity in the neocortex, hippocampus and amygdala ^19,46^. In rodent studies, REM theta coherence between these structures has been linked to fear memory consolidation ^47^, with pharmacological suppression of REM theta disrupting such consolidation ^48^. In human studies, frontal REM theta power was associated with emotional memory performance ^20,49^. Given these results, theta waves might exert some inter-regional coordination that supports reprocessing of distributed memory representations. However, causal evidence establishing this role in human subjects is lacking.

A new approach to causally assess the relationship between sleep oscillations and their function involves targeting stimuli to the oscillation of interest, to effectuate manipulations of the pertaining wave, using automated real-time algorithms. Such closed-loop procedures have for instance been used to influence slow oscillations, spindles and sharp-wave ripples, demonstrating their causal involvement in memory consolidation ^50–54^. For slow oscillations, the dynamic that coordinates information processing during NREM sleep ^55^, it was shown that both the physiological responses and memory effects to auditory stimuli depend not only on the presence of slow oscillations, but also on the phase of the oscillation when the stimulus hits ^56^. Therefore, for this study we developed a novel procedure to precisely phase-target theta waves. The procedure was adapted from a modeling-based closed-loop neurostimulation method (M-CLNS) developed in our lab, that was previously applied to slow oscillations (Fig. 1A) ^57–62^. Given the synchronizing role of REM theta in information processing and its association with emotional memory consolidation, we hypothesize that aligning memory cues to theta waves might enhance emotional memory reprocessing, possibly dependent on the targeted theta phase.

**Figure 1.**
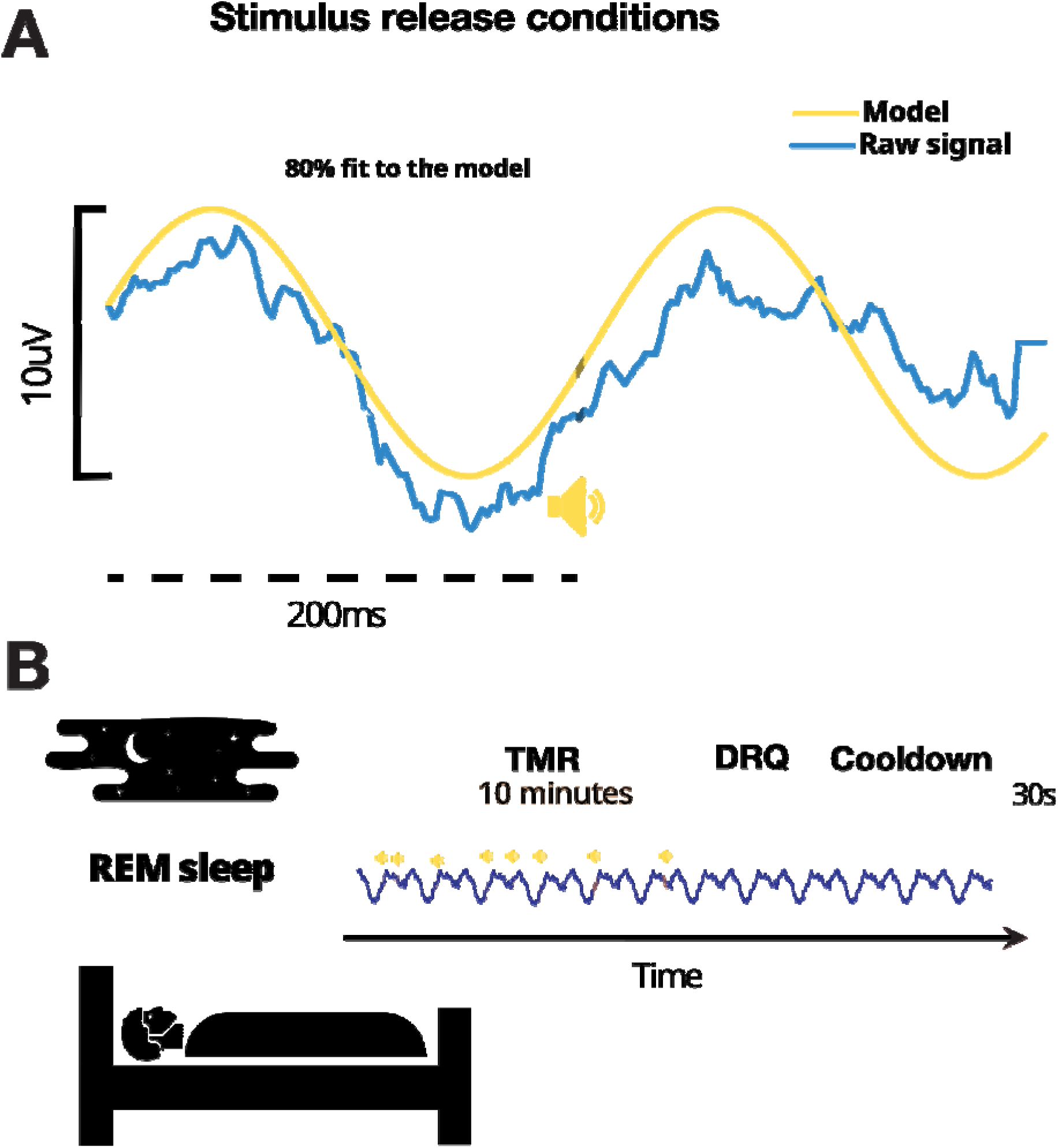
Model-based closed loop neurostimulation protocol. **A**. Sound cue release was determined by a flexible, realtime, sine wave fitting procedure. For each modeling iteration, if (1) the model fit at least 80% with the incoming raw EEG signal and (2) the amplitude of said wave was at least 10uV, a model forecast was used to determine the expected future occurrence of a target theta phase. By doing so, we could anticipate any chosen phase. In the example, a 15ms click was released to coincide with 0° phase (halfway the theta upslope). **B**. For each participant, 10-minute memory reactivation blocks were imposed during REM sleep. During these times the theta phase-targeting algorithm delivered acoustic memory cues related to the prior task according to the group the subject was assigned to (UP, DOWN, SHAM). At the end of a 10-minute TMR block, participants were briefly awakened to answer the dream report questionnaire (DRQ). After that, a refractory period of 45min was ensured before initiating the next TMR block.

Previous REM-TMR studies used pre-existing aversive stimuli to assess declarative memory broadly, confounding episodic and familiarity processes ^42,43^. Therefore, we developed a novel task in which participants were trained to develop novel negative associations with neutral stimuli during the experimental session. This approach allows us to test the effect of REM-TMR on newly formed aversive memories while eliminating potential confounds from pre-established emotional valence.

Theta phase-dependent brain responses to TMR cues were evaluated through evoked potential and time-frequency analyses of the sleep EEG, while effects of TMR cues on emotional memory were assessed through item recognition, valence and arousal assessments, both before and after sleep. We also assessed whether phase-locking to the upswing or down-swing of the theta dynamic influenced the results, akin to what was suggested for SO stimulation during NREM ^56,63,64^. Finally, considering that memory reprocessing might be reflected in dreams, we explored whether theta-phase locked memory reactivation would influence the makeup of dreams.

## Results

### Specific phases of the theta rhythm can be targeted

To implement theta phase-locked TMR during REM sleep, we developed a modified version of our recently introduced modeling-based closed-loop neurostimulation (M-CLNS) algorithm ^57^. This approach builds upon our previous use of the algorithm to phase-target slow oscillations during NREM sleep ^57–60,62^. Following simulation studies, to optimize modeling parameters to theta-phase targeting, we assessed the performance of our M-CLNS method when applied to REM theta oscillations in vivo. Moreover, in the same experiment, we examined the effects of non-arousing acoustic sound stimuli on ongoing neural dynamics (figure S1).

In line with our previous experiments modulating slow oscillations (SOs; 0.5 - 2 Hz) in NREM sleep ^57–59,62^ we were able to accurately model theta dynamics and precisely target sound stimuli to the theta upswing (338 degrees +/- 57.91 degrees). Notably, acoustic stimulation led to a short-lived amplification of the theta dynamic in comparison to sham stimulation. (Methods; Supplementary figure S2).

### Theta-phase targeted TMR

Having validated our theta-targeting approach, we went on to address our main research question. To this aim, we designed a between-subject experiment comprising sixty-one participants separated into three groups: UP (with emotional memory cues placed on the ascending slope of theta oscillations, n = 19), DOWN (emotional memory cues placed on the descending slope of theta oscillations, n = 19) and SHAM (the volume of sound cues is set at 0 dB, n = 23). We opted for targeting UP and DOWN states distinctively in order to effectuate memory reactivation during moments of relatively depolarized (UP) or hyperpolarized (DOWN) cortical states ^65^. During REM sleep each participant underwent up to three 10 minute blocks of theta phase-locked auditory stimulation (Fig. 1A), each followed by a brief interview to obtain dream reports (dream report questionnaire - DRQ; Fig. 1B). Given the prominence of REM sleep theta oscillations in the frontal regions ^20^, the M-CLNS algorithm was applied on the incoming signal recorded from a frontal versus earlobe derivation (Fpz-A1).

For the UP group, the mean effectively targeted phase was 22.85° ± 52.79°(Fig. 2A, left). Similarly, for the DOWN group, we obtained a mean target phase of 127.8° ± 51.05° (Fig. 2A, right).

**Figure 2.**
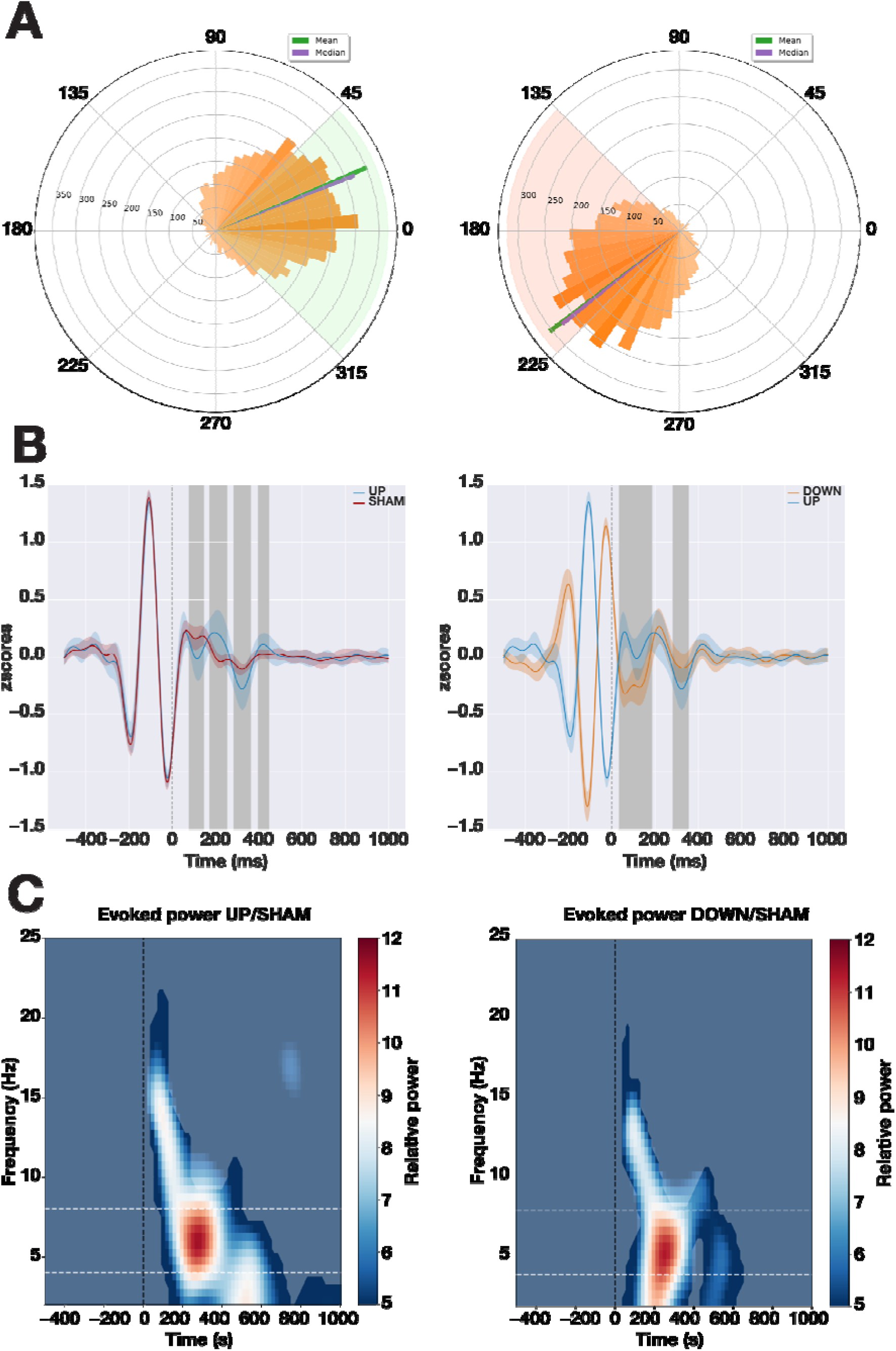
Effects of phase-precise closed loop auditory stimulation on REM theta dynamics. **A**. Circular histograms showing targeting precision for the UP condition (left), at an average of 338° and SD of 57.91°, and the DOWN condition (right), with an average of 220° and SD of 57.91° **B.** Grand-average event-related potentials (ERP) during REM sleep for frontal channel Fz comparing UP stimulation with SHAM stimulation (left), and UP stimulation with DOWN stimulation (right). Error shading indicates standard error of the mean, gray blocks indicate time period of significant difference at cluster level between stimulation conditions (* = P <0.05, *** = P<0.001). **C.** Time-frequency power difference plots for frontal electrode Fz in the UP condition relative to SHAM (left), and DOWN condition relative to SHAM (right). Contours indicate significant differences. Vertical dotted lines indicate stimulus onset (0 s). Horizontal dashed lines indicate the theta range (4-8 Hz).

### REM theta waves are enhanced by modeling-based closed-loop neurostimulation

Stimulus-evoked responses are evident when comparing the event-related potentials (ERPs) in the UP group (Figure 2B, left) to their SHAM counterparts. We observed that sound presentation in the UP group led to a more pronounced theta frequency dynamic compared to SHAM, with a slightly enhanced negativity around 180 ms post-stimulus (permutation test, p < 0.01), a positive wave around 200 ms (p < 0.01), and another negative wave around 350 ms (p < 0.05).

Comparing ERPs for sound presentation between the UP group and the DOWN group (Figure 2B, right), the period prior to t = 0 ms clearly shows the 180° temporal shift in the target phase and therewith in baseline dynamics. Over approximately the first 200 ms after stimulus onset the ERPs still differ significantly (permutation test, p <0.01). This may partially be due to a tendency for the background dynamics to continue. However, over this same period, evoked activity realigns, with a similar enhanced positive deflection observed around 200ms in both the UP and DOWN conditions. This is followed in both conditions by a negative deflection around 350ms, which is larger in the UP group (permutation test, p < 0.05; Figure 2B, right), and again a positive deflection in both groups.

To further assess the effects of acoustic stimulation on ongoing theta dynamics we performed time-frequency analysis (TFA). Relative to SHAM, auditory stimulation in the UP condition evoked an immediate, brief increase in the 10-15 Hz range, followed by an increase in theta power (permutation test, p < 0.01, Figure 2C, left). This theta surge indicates that stimulation-evoked theta power is about twelve times higher in comparison to SHAM. In the DOWN condition, also relative to SHAM, we observe a similar increase in the 10-15 Hz range, followed by a twelve-fold increase in theta power (permutation test, p < 0.01, Fig. 2C, right). There are no significantly different clusters when comparing UP and DOWN TFAs.

### Spatial auditory responses are focused on frontal electrodes

Next, we investigated the effects of theta-phase-locked stimulation across the whole scalp to understand the spatiotemporal profile of the observed enhancement in the amplitude of the theta waves. This was done for the UP, DOWN and SHAM conditions separately. For the UP condition (Fig. 3A), between stimulus onset (t = 0ms) and 300ms afterwards, the response to stimulation appears most prominent in frontal areas, gradually dissipates towards central regions, and is virtually absent in occipital regions. Interestingly, the opposite is observed immediately thereafter, with a stronger occipital effect between 300 and 600 ms. Beyond 600 ms after stimulus onset, the scalp EEG amplitudes does not indicate any significant modulations. Results for the DOWN condition (Fig. 3B) are similar, in that there is a frontally based positive deflection with a main peak around 200ms after stimulus onset followed by, around 400 ms, a frontal negativity and occipital positivity, and only minor modulations after 600 ms. Importantly, both UP and DOWN conditions evoked spatiotemporal patterns of activity significantly different from SHAM throughout the entire period (permutation tests for UP vs SHAM and for DOWN vs SHAM, p < 0.01), but in general not different from each other, with the exception of the first observed peak in the DOWN condition, in the first 157ms after stimulus presentation (permutation test p < 0.01). This difference was not restricted to any particular spatial cluster, but rather was identified across all of the scalp (Fig. 3A and Fig. 3B)

**Figure 3.**
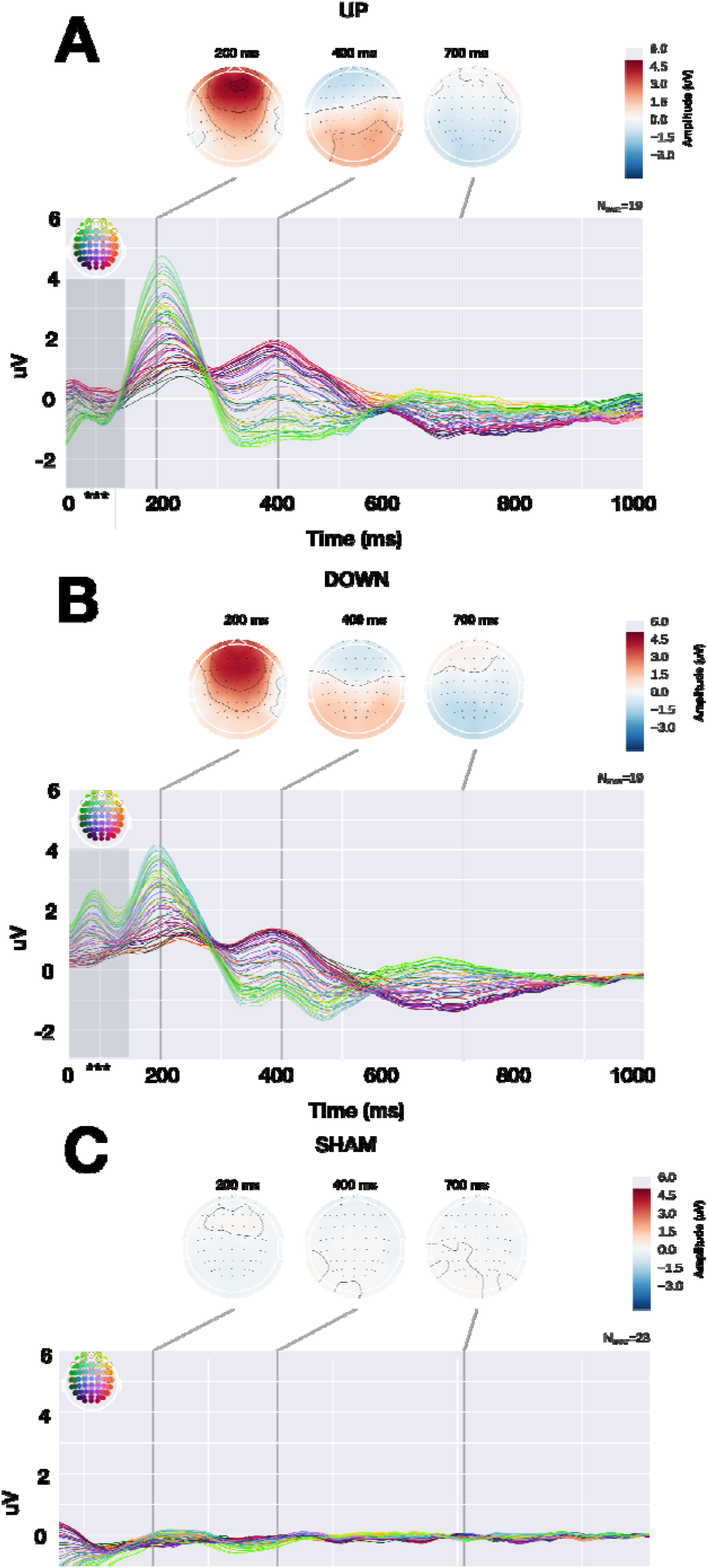
Auditory stimulation in REM theta waves evokes strong frontal activity, regardless of phase. Topographic plots showing effects of presenting memory cues phase-locked to REM sleep theta oscillations in the UP (A), DOWN (B) and SHAM (C) conditions. In the graphs, each line represents one channel, following the color code in the insert displaying electrode positions. Cue onset was at 0 ms. In both the UP and DOWN conditions, stimulation evokes a pronounced effect in the amplitudes of frontal channels (220 ms), that spreads towards the occipital channels (320 – 500 ms) and then dissipates. The SHAM condition (no sound played) indicates that the observed dynamics in (A) and (B) are due to the presentation of sound stimuli, rather than to spontaneous ongoing REM dynamics. Grey bar in panels A and B indicate a significantly different cluster between UP and DOWN stimulation in the first 157 ms after stimulus onset (permutation test, p < 0.01). Both UP and DOWN were different than SHAM for the entire presented analysis window (p < 0.01, permutation test). Topoplots above each graph give an impression of the spatial lay out of signal amplitudes at three time points during the 1000 ms analysis window. Contours in the topoplots indicate electrodes with a similar level of activity.

The SHAM group shows that, in the absence of theta phase-targeted stimulation, spontaneously ongoing theta oscillations quickly dissipate, with no further enhancement or continuity of the waves (Fig. 3C). Alternatively, the lack of clear deflections in the shown time window might reflect either reduced temporal synchronization of neural activity relative to SHAM stimulus onset, decreased theta power, or a combination of these.

These findings suggest that theta phase-locked auditory stimulation evokes a strong frontal component that propagates across the whole scalp.

### Theta-targeted TMR during REM sleep reduces the aversiveness of emotional memories

Next, we proceeded to investigate whether theta phase-targeted TMR during REM sleep, besides enhancing theta dynamics, also influenced emotional memory consolidation.

To this purpose, participants underwent a 2-stage fear conditioning paradigm. Training commenced, for all participants, with a round of habituation (Fig. 4A), during which they listened to two different sounds (short 15ms clicks, one octave apart) and rated the associated valence on a Likert scale. Then, the first stage of fear conditioning began and one of the sounds (CS+; counterbalanced across participants) was associated with a mild shock on the wrist (aversive stimulus, US). After 20 trials in total (10 CS+ and 10 CS-), participants were required to listen to the sounds again and rate their subjective valence on a 9-point Likert scale (1 = very unpleasant, 9 = very pleasant). This ensured that the CS+ had been successfully conditioned. All participants (n = 61) successfully understood the task and provided their ratings, confirming the effectiveness of the conditioning process (CS+ = 4.50 ± 1.08; CS- = 5.92 ± 1.02; mean and standard deviation).

**Figure 4.**
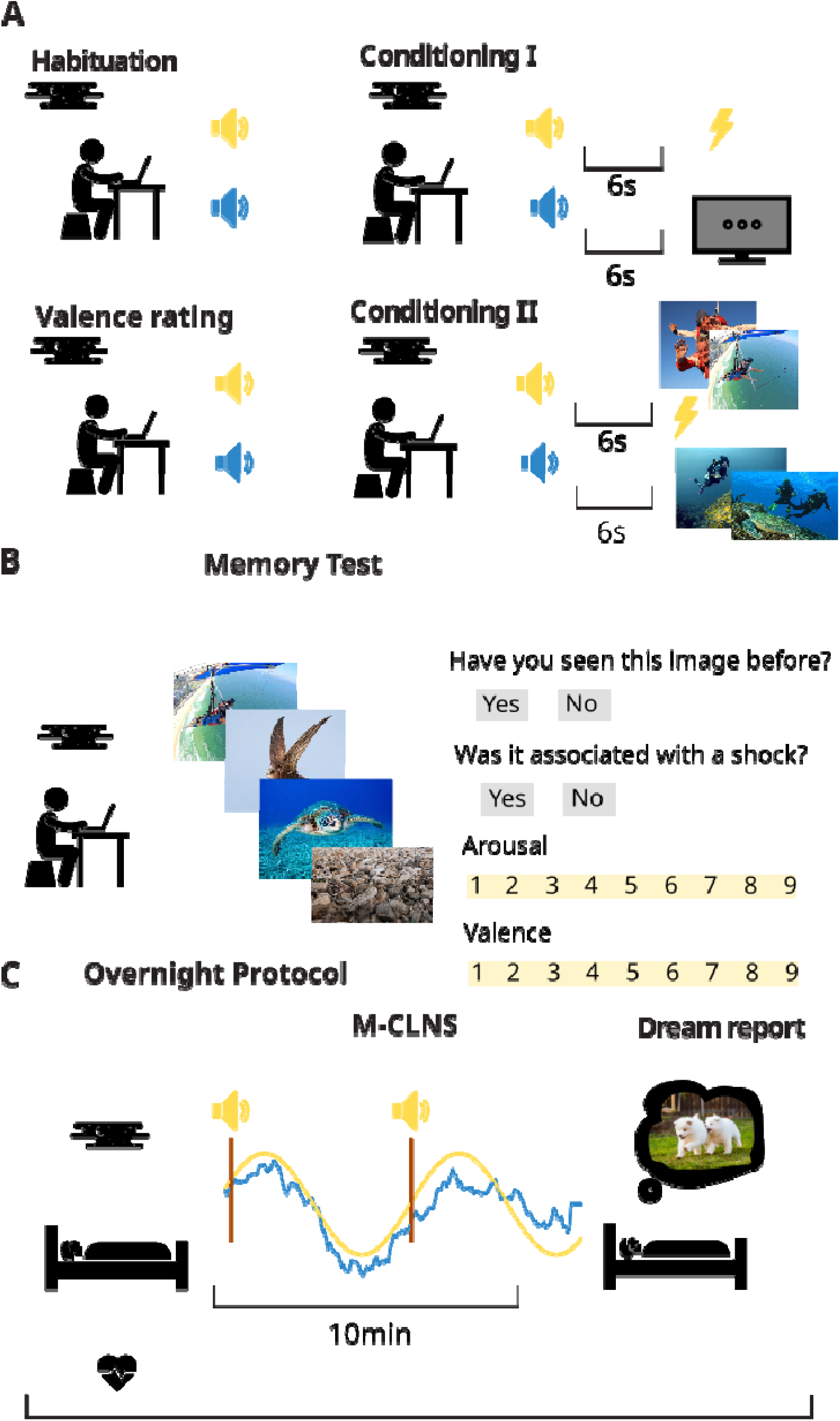
Behavioral protocol. **A.** Participants initially completed a habituation block, where they listened to two different sounds and rated their valence on a Likert scale. Then, one sound (CS+) was paired with an aversive unconditioned stimulus (US), while the other sound (CS-) was presented without any consequence. Participants then rated the valence of each sound again to confirm successful conditioning. In a second conditioning stage, the CS+ and CS- sounds were each paired with images from a specific category (desert or underwater scenes). **B.** After conditioning, participants performed a memory recall task, identifying whether they had seen the presented images before and rating their valence and arousal. **C.** After the immediate recall task, participants went to sleep and underwent theta phase-targeted TMR during REM sleep, in 10-minute blocks, with reexposure to the CS+ sounds to potentially influence emotional memory consolidation. Following each 10-minute TMR block, participants were briefly awakened for a brief interview about their dreaming experience.

In the second stage of the fear conditioning paradigm, the CS+ and CS-were each associated with images belonging to a different category. In other words, the same sound that was associated with the US, would now also be associated with images from one of two categories (desert or underwater images).

After learning, participants were required to perform a memory recollection task (immediate recall; Fig. 4B), during which they were presented with learned items (images presented during the second stage of the fear conditioning paradigm), lures (images belonging to the same category, but not shown during the conditioning paradigm) and foils (images unrelated to the conditioning paradigm). For each item, participants were asked whether they had seen the image before (yes/no) and to rate the valence of the image on a Likert scale ranging from 1 to 9 (1 = totally negative; 9 = totally positive) and arousal on another Likert scale, again from 1 to 9 (1 = totally calm; 9 = totally excited; Fig. 4B).

After completion of the immediate recall task, participants went to sleep and, during the night, underwent theta phase-targeted TMR (Fig. 4C), with re-exposure to the CS+ sound. Upon waking, participants performed the same memory task (Fig. 4B) again (delayed recall), albeit with different learned, lure and foil items.

To assess overnight changes in memory performance across the different groups (UP, DOWN and SHAM), we calculated each participant’s d’ before and after sleep and compared performance on the delayed and immediate recall task, for each group. There was no statistically significant difference between the distribution of the immediate and delayed test for any of the experimental conditions. (Wilcoxon test, p >0.05 for all groups). Moreover, the overnight difference in d’ values did not significantly differ between groups (permutation test, p > 0.05).

Besides recall performance, this task also allowed us to investigate possible overnight changes in memory’s emotional tone, as evidenced by participant’s ratings of each image for valence and arousal. For each participant we calculated the average rating for the images belonging to either the CS+ or CS-. This was done both for immediate and the delayed recall task. Arousal scores were not significantly affected by time or stimulation conditions. However, the overall distribution of participants’ valence scores was significantly affected by overnight cueing in both TMR groups (UP and DOWN; permutation test, p < 0.05, Fig. 5A), with the scores after sleep moving towards a more positive evaluation. This was not the case for the SHAM group (permutation test, p > 0.05, Fig. 5A).

**Figure 5.**
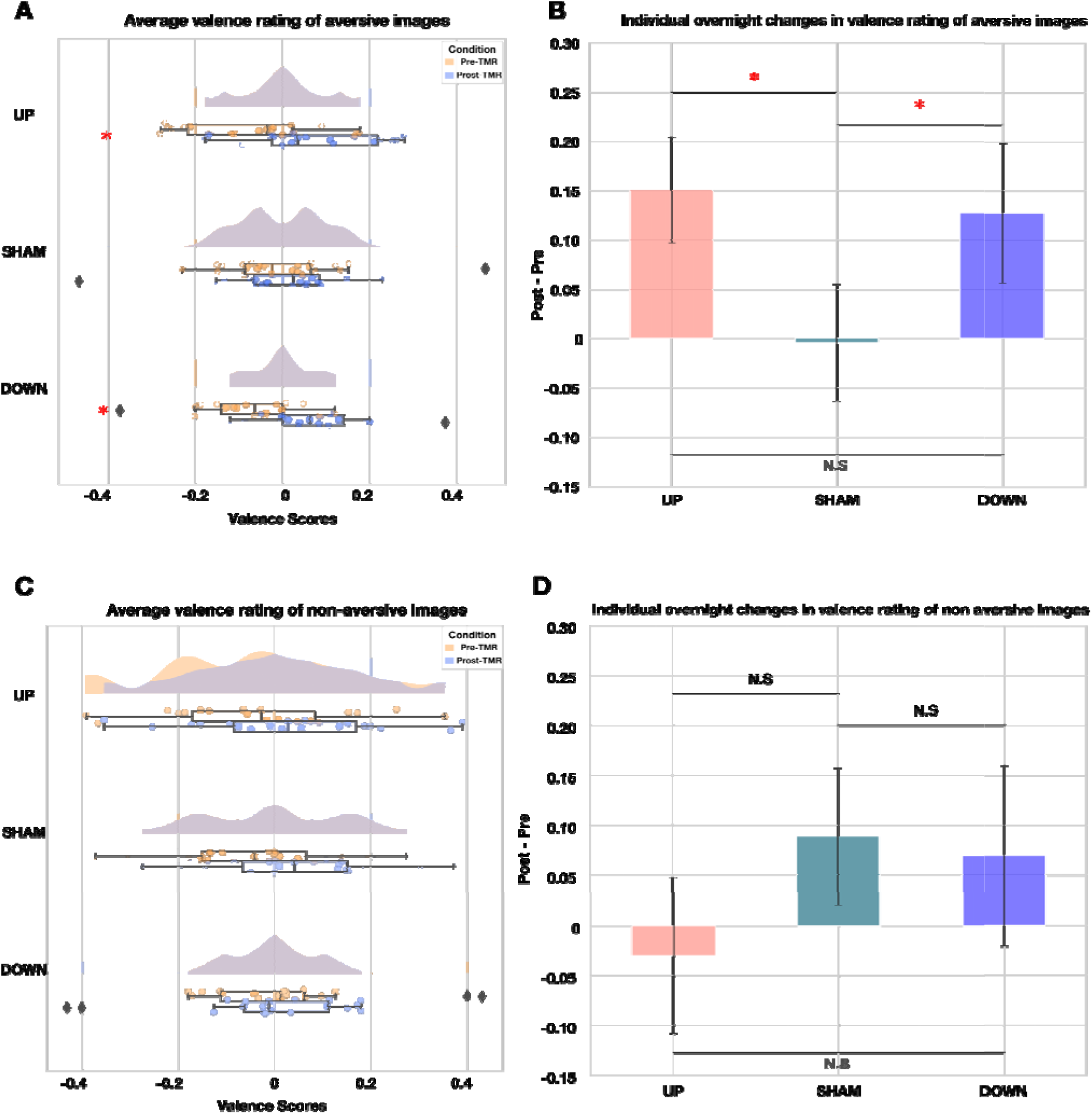
Impact of Targeted Memory Reactivation (TMR) during REM Sleep on Valence Appraisal of Emotional Memories. **A.** Change in Valence Ratings for CS+ Items. Left: The distribution of average valence ratings for CS+ images per participant is displayed before (orange) and after (blue) sleep. Significant differences between these distributions are indicated by asterisks. Both TMR groups (UP and DOWN) showed a significant shift towards more positive evaluations after overnight memory reactivation, compared to the SHAM group. **B.** The difference in valence ratings for CS+ images (postpre) was calculated for each participant, and these differences were averaged across all participants in each group. Both TMR groups (UP and DOWN) demonstrated a significantly greater positive change in valence ratings compared to the SHAM group, indicating that re-exposure to the sound cues during REM sleep led to a more positive appraisal of the aversive images. There was no significant difference between the UP and DOWN groups. **C.** The distribution of average valence ratings for CS-images per participant is shown before (orange) and after (blue) the M-CLNS TMR protocol. There were no significant changes in the evaluations of CS-images overnight for any of the groups. **D.** The individual differences in valence ratings for CS-images (post-pre) were calculated and averaged across participants in each group, revealing no significant differences between any of the groups. This suggests that the observed changes in valence ratings for CS+ items were specific to the re-exposure to the CS+ cues during REM sleep and not a general effect of sleep itself.

Next, we calculated, per participant, the difference in the average valence score per condition (CS+ and CS-) before and after theta phase-targeted TMR and averaged this difference across all participants in the same group (Fig 5B). For the aversive images, we observed a difference between each of the TMR conditions (UP and DOWN) and the SHAM condition (UP vs SHAM, permutation test, p = 0.04, Cohen’s d = 0.44; DOWN vs SHAM, permutation test, p = 0.0071, Cohen’s d = 0.59), indicating that the average change in participants’ evaluations of the images (post vs. pre) was more positive in the TMR groups. Notably, there was no significant difference between the UP and DOWN groups (permutation test, p = 0.731, Cohen’s d = - 0.08; Fig. 5B). Of note, valence ratings for CS-images of all the participants don’t change overnight, either considering the general distributions of valence scores (Fig. 5B, left), or participants’ overnight valence rating differences (Fig. 5B, right). This confirms that, at least in the context of this experiment, more positive valence evaluations are not a global effect of sleep (e.g. through an effect on mood), but rather are related to the re-exposure to the CS+ cues.

Overall, the analysis of memory performance shows that theta-phase locked TMR during REM sleep effectively reduces the aversive tone of fear-encoded memories, without influencing recall.

### Enhanced emotional memory consolidation is not correlated with self-evaluation of dream reports

To investigate whether dreaming activity influenced emotional memory consolidation, we administered a Dream Reports Questionnaire (DRQ) after each block of targeted memory reactivation (TMR). This process involved conducting a brief interview with participants to collect detailed accounts of their dreams and to gather self-rated impressions on several aspects of their dream experiences. Overall, the number of dream reports collected in each TMR group was 33 for UP, 45 for DOWN and 45 for SHAM.

In contrast with the effects observed in behavioral measures (Figure 5), there was no significant difference in the self-reported emotional valence of dream reports between the TMR groups (UP, DOWN, SHAM). This lack of difference was consistent across all evaluated dimensions of the dreams (vividness, p = 0.30; perception of time, p = 0.9; and complexity, p = 0.42). These findings suggest that TMR may directly influence emotional memory consolidation, without significantly altering the subjective qualities of simultaneously occurring dreaming.

## Discussion

In this study, we introduced a novel closed-loop procedure that accurately phase-targets theta oscillations in real-time. By presenting non-arousing acoustic stimuli phase-locked to the upswing or downswing of REM sleep theta waves, we briefly enhanced the amplitude and power of theta waves, compared to sham stimulation. Using this method, we delivered emotional memory cues during REM sleep to reactivate associated memories during periods of theta activity, at specific theta phases. Our results show that theta-targeted TMR promoted overnight reduction of emotional tone, without affecting image recognition, at both tested phases. Overall, these findings provide causal evidence that REM sleep-related memory reactivation specifically dampens the emotional aspects of memories.

Our findings present some similarities with a small number of previous studies that applied emotional memory cues randomly during REM or NREM sleep. One of these found a larger habituation of arousal to negatively charged items for cued compared to uncued items ^42^ when cues were applied during REM sleep, but not during NREM sleep. Two other studies that assessed TMR effects on memory content consolidation ^43,45^ found a benefit for emotional memory cueing only during NREM sleep, not during REM sleep. One of these studies found that REM sleep TMR actually impaired memory^45^, leading the authors to propose REM sleep might be involved in forgetting emotional memories.

Our study design presents several advantages compared to these previous REM-TMR studies. First, while previous studies ^42–45^ assessed TMR effects for naturally aversive stimuli, assessing declarative memory broadly our design isolates episodic memory consolidation, minimizing confounding familiarity effects. Similarly, previous studies used naturally emotional memory cues (e.g. ^42,45^). As such, observed emotional brain responses could be evoked by the cue itself. Conversely, our protocol uses arbitrary (emotionally neutral) memory cues, that have acquired an emotional meaning through conditioning. This ensures that any induced emotional memory processing pertains to the associated memory, rather than the cue itself. In addition, our experiment assessed both memory retention and emotionality scores simultaneously, allowing relative performance on these two modalities across the same items, while previous studies evaluated either memory retention or emotional reactivity. Finally, our task design ensures that per subject and per condition many items are learned and reactivated, allowing robust estimates of memory performance and valence ratings. Moreover, the fact that one TMR cue reactivates an entire category of items, ensures abundant and similar reactivation opportunities for all learned items.

Following a technical innovation, our TMR cues were presented phase-locked to theta waves, showing that such cues temporarily enhance theta oscillations across the whole scalp. (Of note, pilot experiments to develop the theta phase targeting showed that theta enhancement occurs irrespective of whether the sound is a memory cue or an arbitrary sound.) It is possible that therewith theta-linked processes underlying emotional attenuation might also be enhanced, for instance through a temporary, stimulus induced synchronization of the theta dynamic across brain areas involved in memory reprocessing. While further experimentation is needed to understand the role of theta in REM sleep-related emotional processing, theta targeted TMR offers a promising new technique to approach this matter.

Interestingly, studies in rodents are starting to elucidate the neural mechanisms through which REM sleep theta dynamics might lead to reductions in memory’s emotional tone. During REM theta oscillations, transient coupling occurs between the hippocampus, amygdala, and mPFC ^19^. The nature of the coupling between the amygdala and the hippocampus changes over time, transitioning from in-phase to anti-phase theta synchrony with fear extinction learning, marked by reduced freezing behavior in rats ^66–68^. It has been hypothesized that, during reactivation of (hippocampal) memory traces, this anti-phase relation would lead to CA1 input to the amygdala arriving at local theta troughs, which is linked to LTD and depotentiation of CA1–amygdalar synapses ^9^. This could, in turn, lead to an uncoupling of memories’ emotional tone.

More generally (beyond fear conditioning alone), it was found that during rodent REM sleep, the theta phase alignment of spontaneous memory reactivation depends on the novelty of the memory ^67^. Hippocampal CA1 neurons encoding novel environments fire primarily during theta peaks, while those encoding familiar environments burst at theta troughs. This finding has a potential bearing on interpretation of our own findings. Indeed, in our experiment, we examined the effects of REM theta-targeted memory reactivation of newly acquired emotional memories. Our behavioral results revealed that both upslope and downslope memory cueing led to overnight emotional dampening (Fig. 5A and 5B), while neural responses were also similar for both target phases. Possibly, the novelty of the reactivated memories was a more important driver of theta phase alignment than the timing of the memory cue. If so, then, for these newly acquired memories, both up-slope and down-slope targeted TMR cues might have led to theta peak reactivation. This hypothesis could be addressed in future experiments in rodents.

Of note, our theta-phase-targeting algorithm relied on a recently developed approach involving real-time, non-linear sine fitting of raw neuroelectrical signals, combined with model-based signal forecasting ^57,59,60^. The high iteration speed (< 2 ms) of this algorithm enables near-continuous signal predictions, facilitating accurate targeting of stimuli to a large variety of phase and frequency combinations. This adaptive method accommodates temporal variability in neuroelectrical signals, as well as other sources of signal variance (e.g. differences across individuals and sleep-wake stages), almost instantly, without requiring training or parameter customization. This key advantage will enable future studies to address specific functions of specific oscillations, without depending on an adaptation night to set a personalized amplitude threshold ^52^.

## Limitations and Future Directions

Some limitations of our study should be noted. First, given our interest in REM sleep-related emotional memory processing we focused on CS+ cue and sham reactivation during sleep; reactivation of CS-stimuli was outside the scope of the study. The CS-condition was, however, deemed necessary as best practices in fear conditioning require the incorporation of an explicit ‘safe’ condition ^69^. Second, while we compared two theta phases (upslope and downslope) to assess the relevance of phase in memory cueing, considering the similarity of outcomes for both conditions, a definitive assessment of phase relevance will require comparing phase-targeted and random-phase conditions. Third, it is currently not possible to disentangle the memory effects of TMR from those of short-term theta enhancement. Future studies could assess if similar results can be obtained without TMR, by boosting the theta dynamic using meaningless sounds. As a final point, collecting dream reports may have affected sleep quality and task performance, though the impact was likely minimal due to short wake periods (10–25 min). It should also be noted that our analysis of dream reports was limited; more comprehensive evaluations will be conducted in collaboration with dream analysis experts.

## Conclusion

In summary, our study provides strong evidence that memory reprocessing during REM sleep aids the reduction of memories’ emotional tone, with minimal effects on the strength of memories factual content. Moreover, we present a new non-invasive methodology to precisely phase-target and enhance the theta dynamic. Our first use of this technology highlights that REM theta waves may play a crucial role in emotional dampening. Further use of this technology promises to deliver definitive insight into the role of REM sleep related memory reprocessing and may open new avenues for exploring sleep-based interventions for emotional regulation.

## Acknowledgments

This study was supported by the Interdisciplinary Doctorate Agreement Fund of the University of Amsterdam (to UO and LT), by an Amsterdam Brain and Cognition Support Grant (to UO and LT). We thank Christa van der Heijden and Stefan Jongejan for their helpful input and we thank the workshop of the Universiteit van Amsterdam for all their support.

## Methods

| *REAGENT or RESOURCE* | *SOURCE* | *IDENTIFIER* |
| --- | --- | --- |
| **Software and algorithms** |  |  |
| EEGLAB toolbox | https://eeglab.org/ | RRID: SCR_007292 |
| EventIDE software | https://www.okazolab.com | N/A |
| MATLAB | http://www.mathworks.com/products/matlab/ | RRID: SCR_001622 |
| MNE | https://mne.tools/stable/index.html | RRID:SCR_005972 |
| Circular statistics | https://nl.mathworks.com/matlabcentral/fileexchange/10676-circular-statistics-toolbox-directional- | RRID:SCR_016651 |
|  | statis-<br>tics?requestedDomain<br>= |  |
| Python Programming Language | https://www.python.org/ | RRID:SCR_008394 |
| **Other** |  |  |
| Modeling-based closed loop neuro-stimulation | International patent application<br>WO2018156021 | N/A |

### Specific phases of the theta rhythm can be targeted

#### Participants

Twenty-four subjects (20.46 ± 2.06 yr., 19 female, 5 male) from the Faculty of Social and Behavioral Sciences of the University of Amsterdam participated in the experiment for university credit points. Inclusion criteria stipulated that participants self-reported no history of subjective sleep issues or diagnosed sleep disorders, no history of neurological or psychiatric disorder and maintained a regular sleep/wake pattern. Participants were instructed to abstain from alcohol and caffeine on the evening of the study, and were compensated either monetarily or with University of Amsterdam research participation credits. This study was conducted according to the guidelines for ethical principles of the University of Amsterdam.

#### Study design and Procedure

The experiment consisted of a within-subject design, wherein a single experimental night, involving blocks of phase-precise theta stimulation and sham stimulation, was conducted in a dedicated sleep laboratory. Participants arrived at the experimental facilities at 9 PM and were then prepared for polysomnographic recording. They were informed that subtle sounds could be played during the night. A hearing test was performed to assess if they identified the sound stimuli as audible. Next, participants went to bed and lights were turned off at 11PM. When REM sleep was identified (in accordance with AASM guidelines ^70^), a custom-built, sine wave fitting, phase prediction algorithm was deployed in order to target the auditory stimuli to the ascending slope of theta oscillations.

The experimental task was programmed and conducted in EventIDE (Okazolab Ltd, London, United Kingdom) software. Upon deployment of the procedure, two different experimental conditions alternated in blocks of 30s with an interval of 2s between blocks (Figure 1A). For the STIM condition, an auditory stimulus (5ms ‘click’ sound) was played (Figure 1B) and an EEG marker was placed at the exact time of stimulus onset. For the SHAM condition, we followed the same procedure as for STIM, but no sound was played at the predicted target phase. At the detection of an awakening, arousal or whole body movement during the night, the phase prediction algorithm was temporarily halted. Participants were awoken at 8 AM, after a sleep opportunity of 9h. Finally, participants were asked to fill in a general sleep quality questionnaire before departure.

#### Stimulus Material

Designed sound stimuli were 5ms ‘click’ sounds. Every sound was played at 38dB sound pressure, just beneath the arousal threshold during REM sleep (∼42 dB; ^71^). Sounds were delivered over two stereo speakers (GigaWorks T20 Series II, Creative Technology Ltd., Jurong East, Singapore), placed approximately 50 cm from the head of the subject. Before each experimental session, speakers were checked to be delivering at the correct sound pressure levels with a decibel meter (NA-27, Rion Co., Ltd., Kokubunji, Tokyo, Japan).

### Theta phase-targeted TMR during REM sleep -

#### Participants

Sixty-one healthy subjects (34 males, Mean age = 21.05 years, SD = 2.67), recruited among students from the Faculty of Social and Behavioral Sciences of the University of Amsterdam, participated in our experiment for university credit points. Inclusion criteria stipulated that participants self-reported no history of subjective sleep issues or diagnosed sleep disorders, no history of neurological or psychiatric disorder and maintained a regular sleep/wake pattern. On the three days preceding the experiment the subjects were requested to go to bed around 23:00. Moreover, participants were asked to refrain from alcohol, cannabis, or any other drugs from 24 hours preceding the experimental night and to avoid caffeinated drinks as of 14:00.

#### Study design and Procedure

The experiment consisted of a between-subject design, encompassing a 2-stage fear conditioning task and two memory recollection tasks. For all participants, the fear conditioning task was performed in the evening, followed by a memory recollection task (immediate recall). After one experimental night, a second memory task was performed in the morning (delayed recall).

Participants arrived at the sleep laboratory around 7 PM and were prepared for polysomnographic (PSG) recording with hd-EEG. Once the EEG montage was complete, the behavioral procedure started with a habituation block (Fig. 4A), where participants were presented with two sounds (5 times each) that would be used throughout the experiment. After each sound presentation, participants were instructed to rate its valence on a 1-9 Likert scale. Participants that rated any of the sounds as inherently positive or negative were excluded from the task.

After the habituation block, a 2-stage fear conditioning paradigm started. The first part of the paradigm (conditioning I, Fig. 4A) was a classical fear conditioning task, in which a previously neutral stimulus (CS) was associated with an unconditioned stimulus (US). Here, the US was a wrist shock applied to participants’ non-dominant hand using a Digitimer DS7A (Digitimer, Hertfordshire, UK). The intensity of the shocks was self-determined, in a procedure performed together with the experimenters 24h prior to the experimental night ^69^. Participants were thoroughly informed that they had complete control over the intensity of the shocks and could stop the experiment at any given moment. For each participant, one of the two sounds was pseudo-randomly assigned to be coupled to the US (CS+), while the other sound was not (CS-). Participants were exposed to alternating CS+ and CS-trials resulting in 10 of each type. The US was applied at a 80% reinforcement rate of the presentation of the CS+, and was presented 6s after the CS+ sound cue. For both conditions, the total inter stimulus interval was of 7s. After the first conditioning stage was completed, participants were requested to rate each sound on a 9-point Likert scale, where 1 indicated “very unpleasant” and 9 indicated “very pleasant.” Participants who failed to rate the CS+ sound as negative, were excluded from the rest of the experiment.

In the second stage (Conditioning II, Fig. 4A), participants underwent another set of fear conditioning trials, consisting of 50 CS+ (conditioned stimulus) and 50 CS-trials, in random order. During these trials, participants were not only exposed to the US, but also shown images from two different categories (underwater or desert scenes), such that one image category was consistently paired with the CS+, while the other was paired with the CS-. All images had previously been rated as neutral, in terms of both valence and arousal, in a pilot study conducted at the Sleep and Memory Lab (UvA).

In each trial, a CS+ or CS-sound was presented, followed, after 6s, by one image in the corresponding CS+ or CS-category. In the case of CS+, the US was applied concurrently to the image presentation. Each CS+ and CS-was shown randomly 25 times, with an intertrial interval (ITI) of 6 seconds. The image categories were pseudo-randomly assigned to either the CS+ or CS-. Following the first 50 trials, a short break (5 minutes) was issued to avoid a ceiling effect on the US. After the break, a recalibration procedure allowed to adjust the intensity of the wrist shock to the level of “uncomfortable but not painful” ^69^.Hereafter, the task continued for the remaining 50 trials (25 CS+ and 25 CS-trials). After the completion of conditioning, the first memory recollection task (immediate recall) started (Fig. 4B). Here, participants were presented with a total of 100 images that were either learned items (50 images presented during the conditioning paradigm), lures (25 images belonging to the same categories, but not shown during the conditioning paradigm) or foils (25 images unrelated to the conditioning paradigm). For each image, participants were asked whether they had seen the image before (Yes / No). Next, they rated the valence of the image on a Likert scale ranging from 1 to 9 (1 = totally unpleasant; 9 = totally pleasant) and rated arousal, again on a Likert scale from 1 to 9 (1 = totally calm; 9 = totally excited). All the procedures above were programmed and conducted in EventIDE (Okazolab Ltd, London, United Kingdom) software.

After the immediate recall test, participants followed a controlled sleep schedule with lights out at 23:00. For REM theta phase-targeted memory reactivation, participants were subdivided into three different groups: UP (memory cues played during the ascending phase of theta oscillations), DOWN (memory cues played during the descending phase of theta oscillations) and SHAM (no memory cues played, but markers placed in the EEG recording, upon detection of the same conditions as in the UP group).

Theta-targeted memory reactivation occurred in 10-minute blocks, each of which commenced after 5 min of stable, visually inspected REM sleep. At the end of each block, participants were woken up to answer the “dream reports questionnaire” (DRQ) - a short interview, taking ∼7 minutes, to obtain dream content and self-evaluation of emotionality. After the DRQ, participants went back to sleep, and a refractory period of 45 min was ensured before deploying the next TMR block. Per participant a maximum of 3 DRQ blocks were administered to limit sleep interruptions. Stimulation was manually halted whenever participants showed any signs of arousal or transitioning into NREM. After a sleep opportunity of 9h, participants were awoken at 8:00. After waking up, participants filled a final DRQ and thirty minutes after completing the questionnaire (in order to mitigate sleep inertia) they performed the second part (delayed recall) of the memory task, which followed the same procedure as the first one (immediate recall), but with novel learned, lure and foil items, and then were released from the laboratory.

#### Sensory Stimuli

The two-stage fear conditioning task and memory task drew from a collection of 200 images, belonging to two different categories: underwater and desert scenes, which were selected from the Internet. A pilot study was conducted to remove all pictures which were rated outside of the “neutral” range. (see: study design and procedure).

The auditory stimuli consisted of two distinct 15ms sound pulses, each delivered at 38dB sound pressure level, which remained below the arousal threshold for REM sleep (approximately 42dB ^71^). Both sounds were derived from a single ‘click’ sound and digitally modified using Audacity software. The frequency spectra of the two sounds were engineered to differ by one octave while maintaining identical structural characteristics, ensuring they remained clearly distinguishable from each other.

#### EEG recordings and Polysomnography

EEG data was obtained during the night using high-density polysomnography (PSG). Subjects were mounted with a 64-channel WaveGuard EEG cap (ANT, Enschede, The Netherlands) for data acquisition. EEG data was acquired with a 512 Hz sampling rate using a 72-channel Refa DC amplifier (TMS International, Enschede, The Netherlands). Two reference electrodes were placed on the earlobes (location A1 and A2). In addition, two facial electrodes were put near the lateral canthi of the eyes to monitor horizontal electrooculography (HEOG), and another two electrodes were placed above and below the right eye to measure vertical electrooculography (VEOG). Finally, two electrodes were placed under the chin to measure electromyography (EMG). All electrode impedances were kept well below 10 kOhm.

#### Phase-precise targeting procedure

The algorithm used for theta phase targeting was based on our patented modeling-based closed loop neurostimulation method (M-CLNS) (Assembly, method and computer program product for influencing a biological process. WO2018156021 / US 12,310,739 B2, inventors: Talamini, Lucia Maddalena and Korjoukov, Ilia.). Given that theta dynamics are most prominently recorded at frontal regions ^72^, theta phase detection was based on FPz-A1. The algorithm performed real-time, nonlinear sine fitting on the most recent 200 ms of unfiltered EEG signal (iteration time ∼2 ms) and checked the following criteria for stimulus release at each iteration: 1) the fitted sine is in a predefined frequency range of interest (4-8 Hz), 2) the model fit is above an 80% goodness of fit threshold, 3) the predicted target phase occurs within a narrow time window, between 34 and 44 ms, 4) the fitted sine has a minimal amplitude of 10uV. If all criteria were met, sound cues were released to coincide with the upcoming predicted target phase. In order to avoid multiple stimuli being released per theta cycle, a minimal inter-stimulus interval of 250ms was imposed.

#### Preprocessing

Customized scripts and available functions from the EEGLAB toolbox ^73^ were used for all pre-processing steps in Matlab 2016a (Mathworks, Natick, MA). A bandpass filter was applied to the EEG data (0.1 - 100 Hz), as well as a zero-phase Hamming notch filter between 48 - 52 Hz, in order to remove power line electrical artifacts from the recorded signal. EEG channels were then re-referenced to the average signal of the two earlobe electrodes. Epochs were meticulously examined, and those exhibiting signs of arousal or muscle artifacts were excluded from the analysis.

#### Data Analysis

Analyses were conducted in Python 3.10 with the MNE package and custom scripts^74^.

#### Sleep staging

EEG data was re-referenced to the earlobe electrodes, high-pass filtered to 0.1 Hz and low-pass filtered to 35 Hz. EEG channels F3, F4, C3, C4, O1, O2, HEOG, VEOG, EMG were used for offline sleep stage scoring according to AASM criteria ^70^ using the Wonambi package (https://github.com/wonambi-python). The percentage of time spent in each sleep stage was calculated as time in the respective sleep stage over total sleep time (TST).

#### Phase targeting accuracy

Theta phase-targeting performance of the phase prediction algorithm was assessed offline. The EEG signal was filtered between 4 - 8 Hz using a Butterworth bandpass filter order 1. Next, the Hilbert transform was used to obtain the instantaneous phase, and the theta phase distribution across stimulus onsets was assessed. The Circstat toolbox ^75^ was used to perform circular statistics on the phase distributions. Rayleigh test of uniformity was applied to test for uniformity of the distributions and Watson-Williams test for equal means to check for a significant difference in mean accuracy between conditions.

#### Event-related Potentials (ERPs)

To observe the effect of stimulation on theta waves, Fz data was filtered using a Butterworth (first order) high-pass filter of 4 Hz and low-pass filter of 8 Hz and then epoched around stimulus onset (-500 to 1000 ms). Prior to statistical analysis, each trial was normalized by converting the amplitude values to z-scores relative to the pre-stimulus baseline period. Statistical analysis of the ERPs was conducted with non-parametric permutation-based statistical analyses with cluster correction ^76^ using 1000 permutations (false discovery rate corrected significance threshold p < .05).

### Time frequency analysis (TFA)

For time-frequency analysis (TFA), a Butterworth filter of the first order was applied to the signal between 0.5 and 30 Hz. Next, data was epoched around stimulus onset (-500 to 1000 ms). Epochs were then z-scored and averaged per participant, and a Short-time Fast Fourier Transform (SFFT) was applied on the resulting signal to obtain the different frequencies over time. Also here, we applied nonparametric permutation-based statistical analyses with cluster correction^76^. The cluster-alpha parameter was set at alpha = 0.01 and the number of permutations at 1000 (for two-sided testing). Clusters were considered significant at P = 0.025 for two-sided testing.

### Behavioral Analysis

The aim of these analyses was to assess the effects of theta phase-targeted TMR during REM sleep on emotional memory consolidation. To this end, we focused on two key aspects of the task: overnight changes in memory recognition performance, and overnight changes in memory’s affective tone. Memory recognition was quantified through d’, using a signal detection theory approach described by Macmillan and Creelman ^77^. A log linear approach was used to correct for 0 and 1 values by adding 0.5 to the total hits and total false alarms as well as adding a value of 1 to the total signal and noise trials ^78^. Recall performance changes were quantified by calculating the difference in d’ before and after the theta phase-targeted TMR night (post – pre).

Participants’ ratings on the Likert scales for valence and arousal, were z-scored within each group to standardize the data. Affective changes across the theta phase-targeted TMR night were quantified by calculating the difference in the average z-scores before and after sleep (post – pre). Statistical analyses were performed in Python using SciPy, the MNE package, andor custom-built scripts. Comparisons between d-prime differences, valence and arousal scores differences were conducted using permutation-based tests.

## Supplementary Material

**Figure S1:**
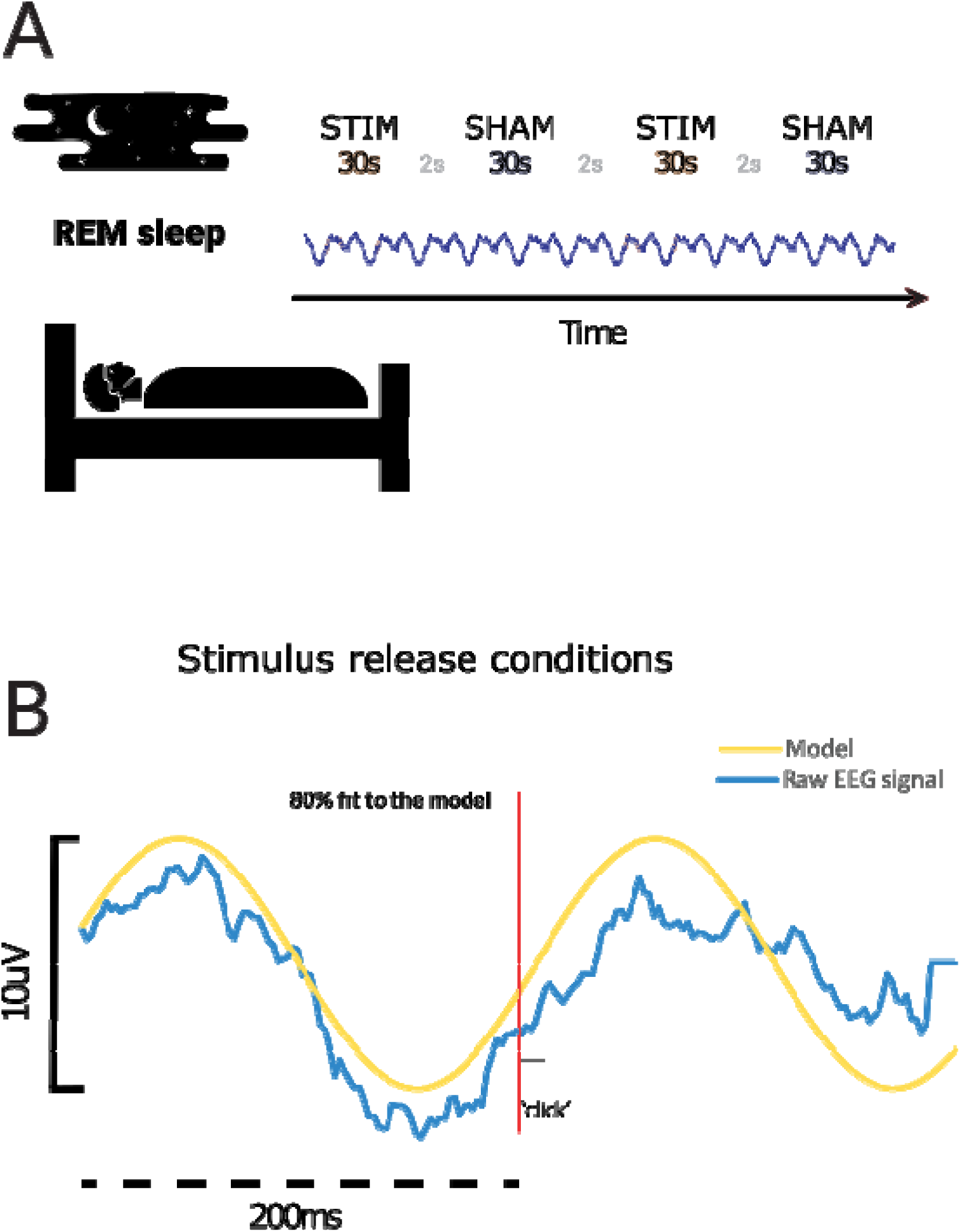
Experimental procedure of targeting of specific phases of theta oscillations. **A.** In a within subject design, during REM sleep of each participant the targeting algorithm was started. In blocks of 30s, both conditions were presented. During STIM, a soft click (38dB) was played, opposed to SHAM when no sound was played. **B.** Sound cue release was determined by a sine wave fitting procedure. In case 1) the model fit at least 80% with the incoming raw eeg signal and 2) the amplitude of said wave was of at least 10uV, we based our forecasting on the model. By doing so, we could anticipate any chosen phase. In this case a 10ms click was released on the 0° phase.

**Figure S2.**
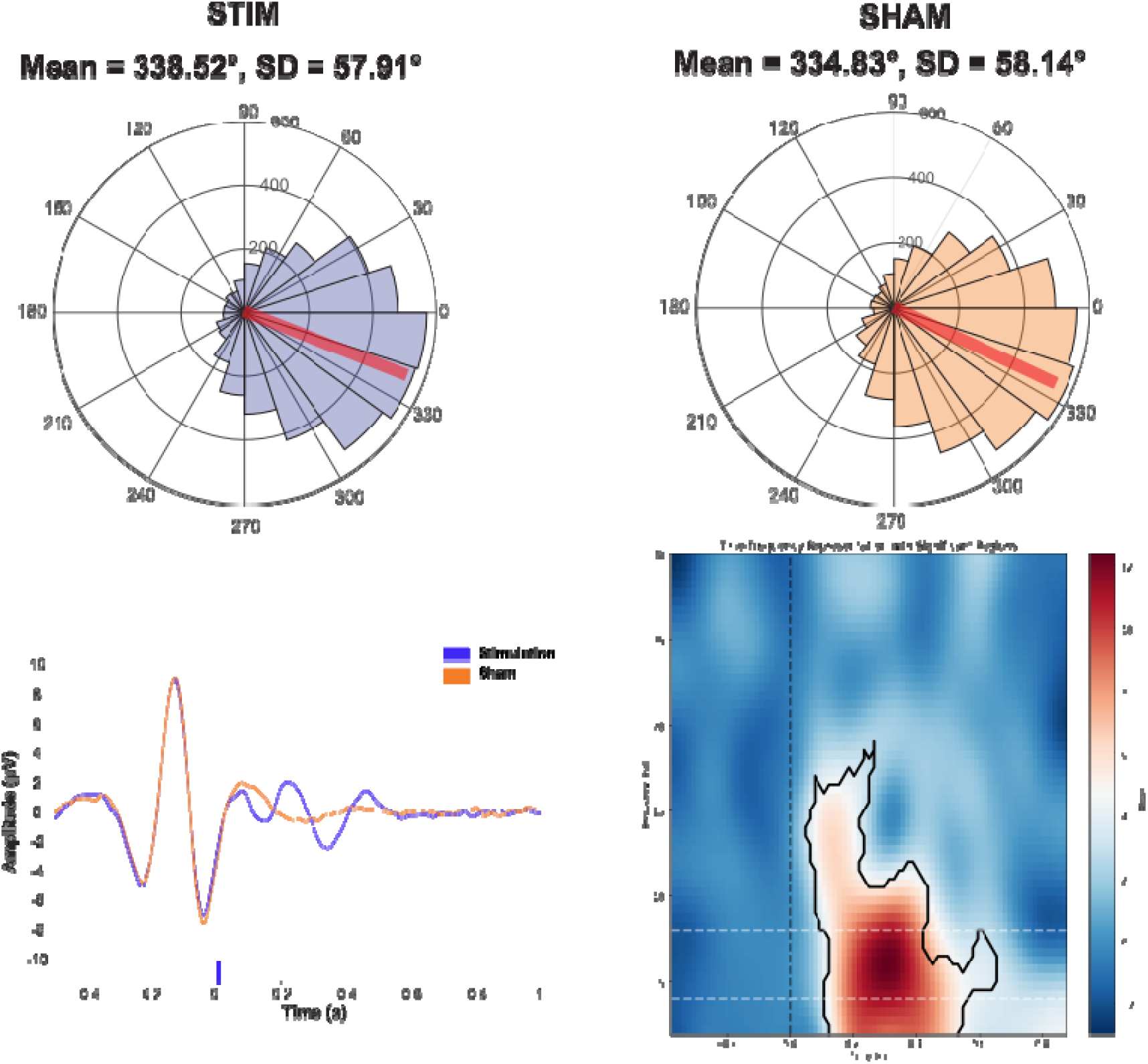
Effects of phase-precise closed loop auditory stimulation in REM theta dynamics. A) Circular histogram showing targeting precision for both stimulation blocks (left) and sham blocks (right). Targeted phase was the ascending phase of the oscillation, with an average of 338° and SD of 57.91°. B) Effects of precise theta auditory stimulation. Left Grand-average event-related potentials (ERP) during sleep for frontal channel Fz. Error shading indicates standard error of the mean, gray blocks indicate time period of significant difference at cluster level between stimulation and sham blocks (* = P <0.05, *** = P<0.001). Right time-frequency power difference plots for frontal electrode Fz. Similar to the ERP, contours indicate significant differences. Vertical dotted lines indicate stimulus onset (0 s). Horizontal dashed lines indicate the theta range (4-8 Hz).

